# Cotranslational folding and assembly of the dimeric *E. coli* inner membrane protein EmrE

**DOI:** 10.1101/2022.04.02.486806

**Authors:** Daphne Mermans, Felix Nicolaus, Klara Fleisch, Gunnar von Heijne

## Abstract

In recent years, it has become clear that many homo- and heterodimeric cytoplasmic proteins in both prokaryotic and eukaryotic cells start to dimerize cotranslationally, *i*.*e*., while at least one of the two chains is still attached to the ribosome. Whether this is possible also for integral membrane proteins is unknown, however. Here, we apply Force Profile Analysis (FPA) – a method where a translational arrest peptide (AP) engineered into the polypeptide chain is used to detect force generated on the nascent chain during membrane insertion – to demonstrate cotranslational interactions between a fully membrane-inserted monomer and a nascent, ribosome-tethered monomer of the *E. coli* inner membrane protein EmrE. Similar cotranslational interactions are also seen when the two monomers are fused into a single polypeptide. Further, we uncover an apparent intrachain interaction between E^14^ in TMH1 and S^64^ in TMH3 that forms at a precise nascent chain length during cotranslational membrane insertion of an EmrE monomer. Like soluble proteins, inner membrane proteins can thus both start to fold and start to dimerize during the cotranslational membrane-insertion process.

**Significance statement:** Many water-soluble proteins are known to fold and even dimerize cotranslationally, *i*.*e*., when still attached to the ribosome. Here, we show that an *E. coli* inner membrane protein can also start to fold and dimerize cotranslationally, establishing the generality of these cotranslational maturation processes.

## Introduction

It is becoming increasingly clear that many, if not most, cytoplasmic proteins start to fold cotranslationally, *i*.*e*., while the growing nascent polypeptide chain is still attached to the ribosome. Such early folding events range from the formation of elements of secondary structure and small protein domains already within the ribosome exit tunnel to folding of larger domains just outside the exit tunnel, with or without the help of chaperones (1, 2). Ribosome profiling experiments in both prokaryotic and eukaryotic cells have further shown that many homo- and heterodimeric cytoplasmic proteins can start to dimerize cotranslationally while one or even both monomers are still attached to the ribosome (3). Cotranslational folding and assembly of soluble proteins thus seem to be common phenomena; however, whether this is true also for integral membrane proteins remains unclear. Individual domains in multi-domain membrane proteins such as the Cystic Fibrosis Transmembrane Conductance Regulator (CFTR) or the *E. coli* inner membrane protein GlpG fold mainly cotranslationally (4-7), but to what extent individual transmembrane helices (TMHs) can interact during translocon-mediated membrane insertion and whether ribosome-attached, nascent integral membrane proteins can start to dimerize with already folded partner proteins are still open questions.

To address these issues, we decided to perform an in-depth Force Profile Analysis (FPA) of the cotranslational membrane insertion process of the small multidrug-resistance protein EmrE from *E. coli*. EmrE has four TMHs and is a dual-topology protein, *i*.*e*., the monomers integrate into the inner membrane in a 50-50 mixture of N_in_-C_in_ and N_out_-C_out_ topologies; oppositely oriented monomers then assemble into antiparallel dimers (8, 9). A recent FPA analysis of EmrE suggested that there may be long-range cotranslational interactions between a conserved Glu residue (E^14^) in the middle of transmembrane helix 1 (TMH1) and unidentified residues in TMH2 and TMH3 during membrane insertion (7), and hence that the monomer might start to fold cotranslationally. Further, given the extensive inter-subunit packing interactions between the two monomers in the EmrE dimer (10), we speculated that cotranslational dimerization of EmrE might be possible to observe by FPA.

FPA takes advantage of so-called translational arrest peptides (APs) – short stretches of polypeptide that bind with high affinity in the upper reaches of the ribosome exit tunnel and thereby arrest translation at a specific codon in the mRNA (11). The translational arrest can be overcome if a strong enough pulling force is exerted on the AP, essentially pulling it out of its binding site in the exit tunnel (12-16). APs can be employed as sensitive ‘molecular force sensors’ to report on various cotranslational events such as protein folding (17, 18), protein translocation (19, 20), and membrane protein integration (7, 13).

Using FPA, we have now identified a residue in EmrE TMH3 (S^64^) that appears to form a specific interaction with E^14^ in TMH1 at a precise point during the cotranslational membrane insertion process. We also show that TMH4 in one EmrE monomer can interact cotranslationally with TMH4 in a second, already fully membrane-inserted monomer and, similarly, that the TMH4 transmembrane helices in a construct where two EmrE monomers have been fused into one polypeptide can interact cotranslationally. Cotranslational folding and dimerization events are thus not restricted to soluble proteins but can also be observed in integral membrane proteins.

## Results

### Force Profile Analysis

FPA is based on the ability of APs to bind in the upper parts of the ribosome exit tunnel and thereby pause translation when their last codon is in the ribosomal A-site (11). The duration of an AP-induced pause is reduced in proportion to pulling forces exerted on the nascent chain (14, 21), *i*.*e*., APs can act as force sensors, and can be tuned by mutation to react to different force levels (22). In an FPA experiment, a series of constructs is made in which a force-generating sequence element (*e*.*g*., a TMH) is placed an increasing number of residues away from an AP (here, we use the AP from *E. coli* SecM (23)), which in turn is followed by a C-terminal tail, Fig. 1a (construct lengths are denoted by *N*, the number of residues from the start of the protein to the end of the AP). In constructs where a TMH engages in an interaction that generates a strong enough pulling force *F* on the nascent chain at the point when the ribosome reaches the last codon of the AP, pausing will be prevented and mostly full-length protein will be produced during a short pulse with [^35^S]-Met, Fig. 1b (left). In contrast, in constructs where little force is exerted on the AP, pausing will be efficient and more of the shorter, arrested form of the protein will be produced, Fig. 1b (right). The fraction full-length protein produced, *f*_*FL*_ = *I*_*FL*_/(*I*_*FL*_+*I*_*A*_), where *I*_*FL*_ and *I*_*A*_ are the intensities of the bands representing the full-length (*FL*) and arrested (*A*) species on an SDS-PAGE gel, Fig. 1c (see Supplementary Fig. S1 for SDS-PAGE gels of all constructs), can therefore be used as a proxy for *F* in a given construct (21, 24, 25). A plot of *f*_*FL*_ vs. *N* – a force profile (FP) – thus can provide a detailed picture of the cotranslational process in question, as reflected in the variation in the force exerted on the nascent chain during translation, Fig. 1d (see Supplementary Table S1 for numerical *f*_*FL*_ values for all constructs). FPs can be recorded with up to single-residue resolution by increasing *N* in steps of one residue (corresponding to a lengthening of the nascent chain by ∼3Å).

**Figure 1.**
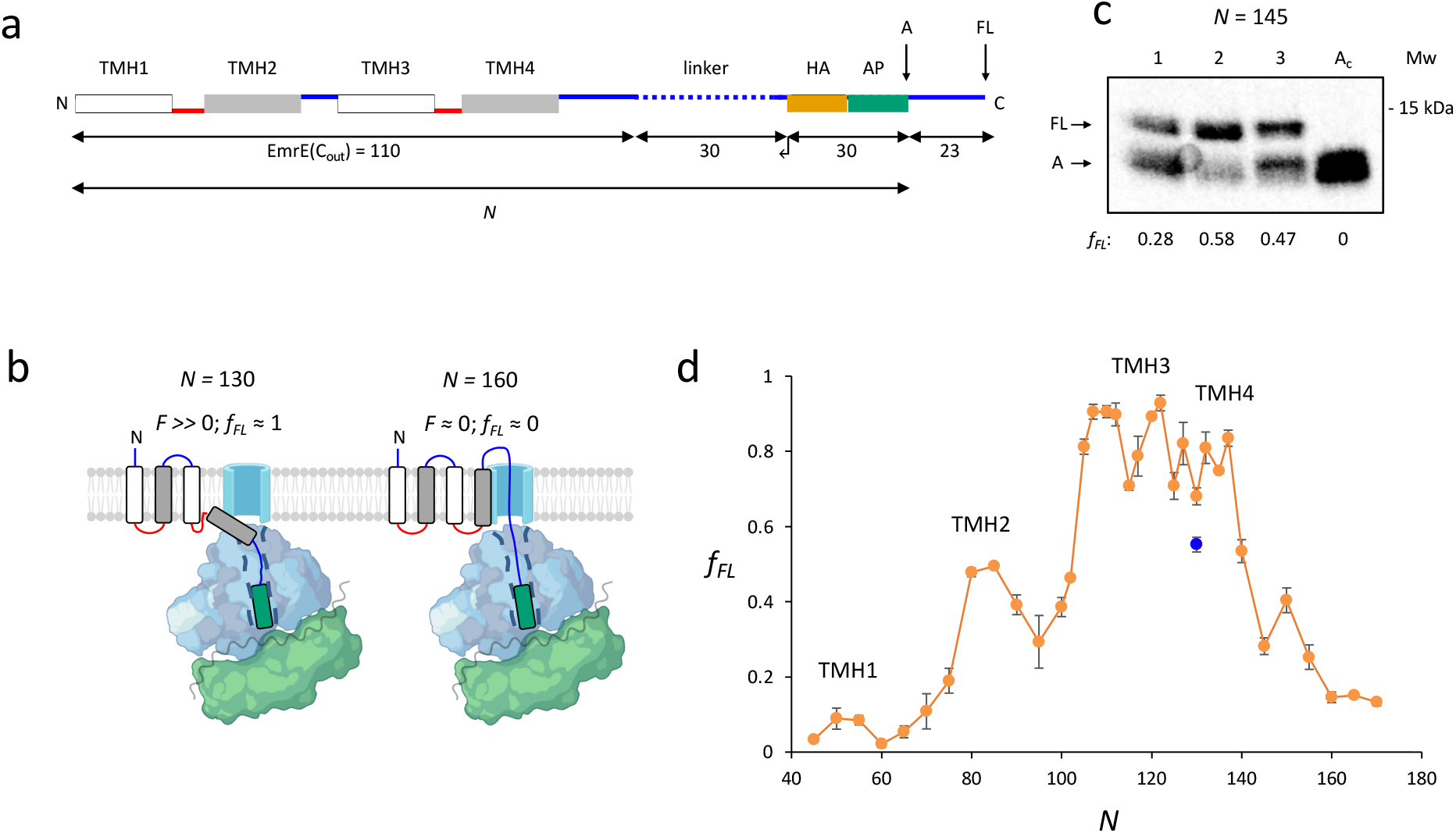
The force profile assay. (a) Basic EmrE(C_out_) construct. To obtain a FP, EmrE(C_out_) is shortened from the C-terminal end of the LepB-derived linker (dotted), as indicated by the arrow. Cytoplasmic (red) and periplasmic (blue) loops, and lengths of full-length EmrE(C_out_), LepB-derived linker, hemagglutinin tag and arrest peptide (HA+AP), and C-terminal tail are indicated. Construct lengths are denoted by *N*, the number of residues between the N-terminal end of EmrE(C_out_) and the C-terminal end of the AP. Since the 30-residue HA+AP segment is constant in all constructs, the FP reflects nascent chain interactions occurring mainly outside the ribosome exit tunnel. (b) At construct length *N* = 130 residues, TMH4 is starting to integrate into the membrane, generating a high pulling force on the nascent chain. At *N* = 160 residues, TMH4 has finished integrating into the membrane and generates little pulling force. (c) SDS-PAGE gel showing [^35^S]-Met labelled and immunoprecipitated EmrE(C_out_) [*N* = 145] (lane 1), EmrE(C_out_) [*N* = 145] produced in the presence of co-expressed EmrE(C_in_) (lane 2), and EmrE(C_out_) [*N* = 145] produced in the presence of co-expressed EmrE(C_in_;G^90^P+G^97^P) (lane 3). Control construct *A*_*C*_ has a stop codon replacing the last Pro codon in the AP in EmrE(C_out_) [*N* = 145] (lane 4). Average *f*_*FL*_ values are indicated. (d) FP for EmrE(C_out_) (orange; adapted from (7)). The peaks corresponding to the membrane insertion of TMH1-4 are indicated (7). Error bars indicate SEM values. The *f*_*FL*_ value for construct EmrE(C_out_;E^14^L) [*N* = 130] (blue data point) is significantly different from the corresponding value for EmrE(C_out_) [*N* = 130] (*p* = 0.002, two-sided t-test). Sequences for all constructs used in this study are listed in Supplementary File S1 and all *f*_*FL*_ values in Supplementary Table S1.

### Cotranslational interactions between TMH1 and TMH3 in the EmrE monomer

In our recent study of the cotranslational membrane insertion of EmrE(C_out_) (7) – a mutant version of EmrE that inserts only with N_out_-C_out_ orientation (9) – we found that mutation of the key functional residue E^14^ in TMH1 to Leu gave rise to significant changes in the FP at three specific nascent chain lengths: *N* = 85, 115, and 130 residues. We decided to focus on the *N* = 130 construct, Fig. 1d, as mutation of E^14^ to a hydrophobic (Leu, Ala) but not polar or charged (Gln, Asp) residue led to a significant reduction in the *f*_*FL*_ value at *N* = 130 (7), suggesting the formation of a polar interaction between E^14^ and some other residue in the protein when the nascent chain reaches an overall length of *N* = 130 residues. At this chain length, TMH4 (residues 88-103) is about to begin inserting into the membrane and TMH3 (residues 56-78) should just have reached its membrane-spanning disposition, Fig. 1b (left). In the EmrE dimer, TMH1 is sandwiched between TMH2 and TMH3 in each monomer, Fig 2a. We therefore considered the polar residues in TMH3 (Y^60^, W^63^, S^64^, W^76^; Fig. 2a) as the best candidates for making a specific interaction with E^14^ at *N* = 130 residues. These four residues were individually mutated to Ala, both in EmrE(C_out_) and EmrE(C_out_;E^14^L).

**Figure 2.**
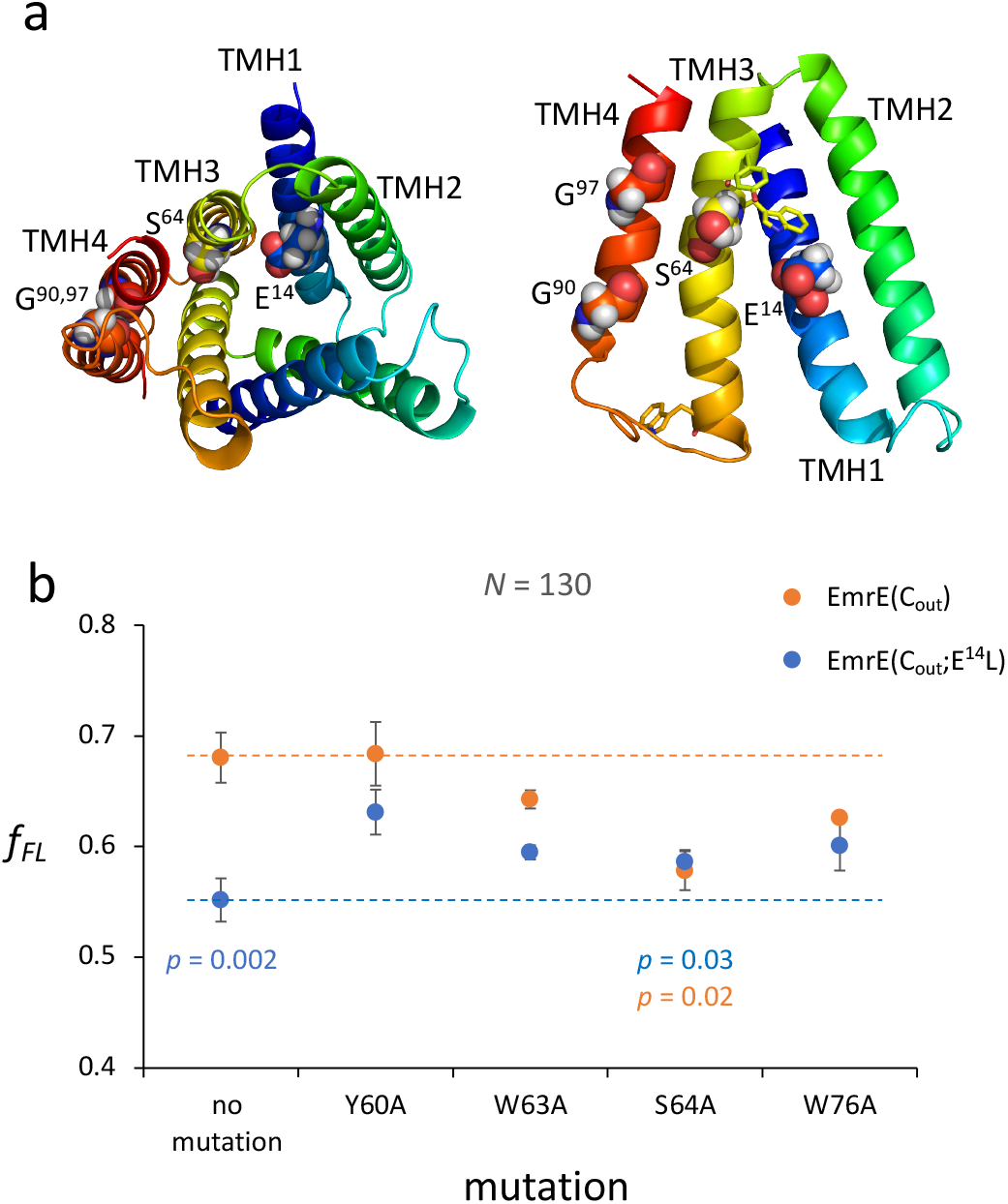
Identification of cotranslationally interacting residues in EmrE(C_out_). (a) The EmrE dimer (left) and one monomer (right) (10) (PDB 7MH6). E^14^, S^64^, G^90^, and G^97^ are shown in spacefill and the other residues mutated in TMH3 are indicated in stick representation. (b) *f*_*FL*_ values for mutant versions of EmrE(C_out_) (orange data points) and EmrE(C_out_;E^14^L) (blue data points) at *N* = 130 residues. Error bars indicate SEM values (*n* ≥ 3, see Supplementary Table S1). The *p*-values for EmrE(C_out_;E^14^L), EmrE(C_out_;S^64^A), and EmrE(C_out_;E^14^L+S^64^A) compared to EmrE(C_out_) are shown. *p*-values were calculated by a two-sided t-test.

In general, in the absence of specific interactions between TMH3 and upstream TMHs, polar-to-hydrophobic mutations in TMH3 are expected to increase the pulling force generated during its membrane insertion (13), leading to increases in *f*_*FL*_. As seen in Fig. 2b, this is indeed seen at *N* = 130 residues for all four mutations when made in the EmrE(C_out_;E^14^L) background (blue data points). In contrast, three of the four mutations tend to decrease *f*_*FL*_ when made in the EmrE(C_out_) background (orange data points). The strongest reduction is seen for S^64^A, which reduces *f*_*FL*_ at *N* = 130 residues to the same extent as does the E^14^L mutation in TMH1 (from 0.68 to 0.58). The double mutation E^14^L + S^64^A has no further effect on *f*_*FL*_. These results suggest that a stabilizing interaction is formed between E^14^ in TMH1 and S^64^ in TMH3 at *N* = 130 residues. Indeed, assuming that TMH1-TMH3 in the monomer can adopt a structure similar to that seen in the dimer, S^64^ is well placed to interact with E^14^, as seen in Fig. 2a.

### Cotranslational assembly of the EmrE dimer

Many soluble cytoplasmic proteins can form both homo- and heterodimers while one of the partner proteins is still being translated (3). Here, we wanted to ascertain whether this is possible also for EmrE that assembles into an anti-parallel 4+4 TMH homodimer in the inner membrane (10, 26, 27), Fig. 2a.

It has been shown that efficient dimerization of EmrE depends critically on a tight interaction between the TMH4 helices in the two monomers (28), and we therefore focused our attention on the part of the FP that reports on the membrane insertion of TMH4, *i*.*e*., *N* ≈ 130-170 residues (*c*.*f*., Fig 1d). In a first set of experiments we recorded a FP for EmrE(C_out_) while co-expressing an oppositely oriented EmrE(C_in_) version that is known to dimerize efficiently with EmrE(C_out_) (9, 29, 30), Fig. 3a. Indeed, as shown in Fig. 3b, the presence of EmrE(C_in_) causes a shoulder in the EmrE(C_out_) FP in the region N ≈ 140-150 residues (magenta data points) where *f*_*FL*_ is significantly increased compared to the EmrE(C_out_) FP (orange data points), suggesting a cotranslational interaction between TMH4 in the nascent EmrE(C_out_) subunit and the already synthesized EmrE(C_in_). We further recorded a FP for EmrE(C_out_) with co-expression of a version of EmrE(C_in_) carrying Gly→Pro mutations in positions 90 and 97 in TMH4 (see Fig. 2a) that are known to destabilize the heterodimer (28, 31), Fig. 3c. Indeed, the EmrE(C_out_) FP obtained while co-expressing EmrE(C_in_;G^90^P+G^97^P) (light blue data points) was closer to the original EmrE(C_out_) FP obtained in the absence of co-expressed EmrE(C_in_).

**Figure 3.**
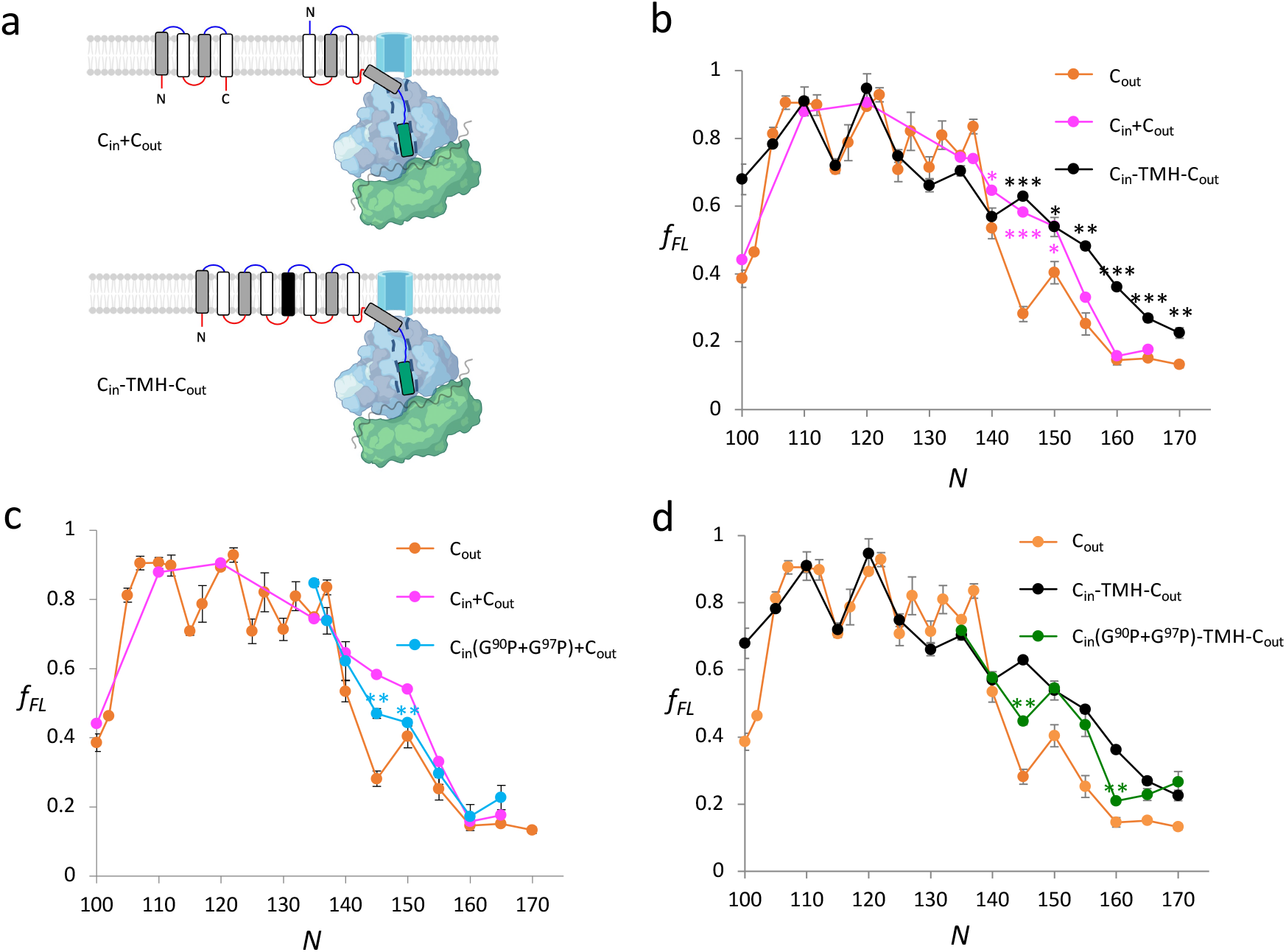
Cotranslational assembly of the EmrE dimer. (a) FPs for EmrE(C_out_) were obtained while co-expressing EmrE(C_in_) (upper panel) and for an EmrE(C_in_)-TMH-EmrE(C_out_) fusion construct in which a TMH of composition 7L/12A (black) was included to maintain the opposite orientations of the EmrE(C_in_) and EmrE(C_out_) moieties (lower panel). *N* values were counted from the N-terminal residue of EmrE(C_out_) in both cases. (b) FPs for EmrE(C_out_) (orange data points), EmrE(C_out_) with co-expressed EmrE(C_in_) (magenta data points), and fused EmrE(C_in_)-TMH-EmrE(C_out_) (black data points). *p*-values were calculated by a two-sided t-test comparing the two latter sets of data points to those of EmrE(C_out_) (*p* < 0.05: *; *p* ≤ .01: **; *p* ≤ 0.001: ***). (c) FPs for EmrE(C_out_) (orange), EmrE(C_out_) with co-expressed EmrE(C_in_) (magenta), and EmrE(C_out_) with co-expressed EmrE(C_in_;G^90^P+G^97^P) (light blue). *p*-values were calculated comparing the latter set of data points to those of EmrE(C_out_) with co-expressed EmrE(C_in_). (d) FPs for EmrE(C_out_) (orange), fused EmrE(C_in_)-TMH-EmrE(C_out_) (black), and fused EmrE(C_in_; G^90^P+G^97^P)-TMH-EmrE(C_out_) (green). *p*-values were calculated comparing the latter set of data points to those of EmrE(C_in_)-TMH-EmrE(C_out_). In all cases, the FPs are for the EmrE(C_out_) subunit. Error bars indicate SEM values (*n* ≥ 3).

To ascertain whether the cotranslational interaction requires that EmrE(C_in_) is expressed from the same mRNA as EmrE(C_out_) (*i*.*e*., in *cis*), we modified the pET-Duet-1 plasmid used to co-express EmrE(C_in_) with EmrE(C_out_). pET-Duet-1 has two T7 promoters but no intervening transcriptional terminator, and we therefore recorded two additional FPs, one in which the second T7 promoter, located upstream of the EmrE(C_out_) ORF, was deleted (ΔT7-2) and one in which the strong, tri-partite tZ terminator (32) was inserted between the EmrE(C_in_) ORF and the second T7 promoter, Supplementary Fig. S2. The two FPs were essentially identical to each other and to the original EmrE(C_in_)+EmrE(C_out_) FP. Hence, the cotranslational interaction between EmrE(C_in_) and EmrE(C_out_) is seen regardless of whether the two subunits are expressed in *cis* or in *trans*.

Finally, we recorded a FP for a fusion construct between EmrE(C_in_) and EmrE(C_out_) (with an extra TMH inserted between EmrE(C_in_) and EmrE(C_out_) in order to maintain their anti-parallel orientations in the membrane), Fig. 3a. The presence of EmrE(C_in_), now covalently fused to the N-terminus of EmrE(C_out_), caused an even more conspicuous shoulder in the EmrE(C_out_) FP, Fig. 3b (black data points). Again, introduction of the G^90^P+G^97^P mutations in the fused EmrE(C_in_) part partially reverted this effect, Fig. 3d (green data points).

We conclude that the presence of EmrE(C_in_) during expression of EmrE(C_out_) gives rise to a clear increase in the *f*_*FL*_ values in the *N* ≈ 140-150 residues region of the FP (and in an even longer region when the two subunits are fused together). The G^90^P+G^97^P mutation in EmrE(C_in_) TMH4 reduces this effect. According to our earlier work, EmrE(C_out_) TMH4 starts to insert into the membrane at *N* ≈ 132 residues and stops generating a pulling force on the nascent chain at *N* ≈ 150 residues (7), *i*.*e*., the cotranslational interaction seen between EmrE(C_in_) and EmrE(C_out_) corresponds to the last steps in the membrane insertion of TMH4. The cotranslational interaction seen in the FP recorded for the fused subunits extends beyond this point, suggesting that other, presumably weaker, interactions between the two subunits also come into play in this case.

## Discussion

Thanks to the high resolution and sensitivity of FPA, we have been able to identify the cotranslational formation of what appears to be a specific interaction between two EmrE residues – E^14^ in TMH1 and S^64^ in TMH3 – at the point when TMH3 is just completing its insertion into the inner membrane. The interaction is seen as a small increase in *f*_*FL*_ at *N* = 130 residues, which disappears when either E^14^ or S^64^ is mutated to a non-polar residue. Thus, TMH1 and TMH3 appear to interact cotranslationally within the context of the SecYEG translocon. We have also found that the EmrE anti-parallel dimer can start to assemble in the inner membrane while one of the two monomers is still attached to the ribosome (albeit by an artificial C-terminal tether). The first clear signal of dimerization is seen at *N* ≈ 145 residues (at which point the C-terminal end of TMH4 is ∼40 residues from the PTC), corresponding to a situation where TMH4 in the EmrE(C_out_) monomer is close to being fully inserted into the membrane and must still be in, or in the immediate vicinity of, the SecYEG translocon. Thus, EmrE(C_in_) monomers must have access to the SecYEG translocon at this point, which may not be so surprising in the case of the EmrE(C_in_)-TMH-EmrE(C_out_) fusion construct but is quite remarkable in the case of co-expressed EmrE(C_in_) and EmrE(C_out_). More generally, our results show that, just like cytoplasmic proteins (3), inner membrane proteins can undergo cotranslational folding and dimerization, adding a new level of complexity to the basic two-stage model for membrane protein folding (33, 34).

## Materials and methods

| *Reagent type (species) or resource* | *Designation* | *Source or reference* | *Identifiers* | *Additional information* |
| --- | --- | --- | --- | --- |
| Strain, strain background (*Escherichia coli*) | BL21(DE3) | Sigma-Aldrich | CMC0016 | Electrocompetent cells |
| Other | Rifampicin | Sigma-Aldrich | R3501 | Used for inhibition of bacterial RNA polymerase during expression |
| Other | Protein G-Agarose | Roche | 11243233001 | Resin used for immunoprecipitation |
| Antibody | Anti-HA.11 Epitope Tag Antibody (mouse monoclonal) IgG | BioLegend | Cat# 901533 | Used for immunoprecipitation (1 µl of 1 mg/mL, diluted 1:820) |
| Recombinant DNA reagent | pET-Duet-1 (plasmid) | Novagen | Cat # 71146 | Expression plasmid |
| Commercial assay, kit | GeneJET Plasmid miniprep kit | Thermo Fisher Scientific<br>RRID:SCR_008452 | Cat no. 0502 | Used to purify plasmids |
| Chemical compound, drug | <sup>35</sup>S Methionine | Perkin-Elmer | Cat no. NEG009T001MC | [<sup>35</sup>S]-Met is incorporated into the protein during *in vitro* and *in vivo* translation and aids detection by phosphor-imaging. |
| Software, algorithm | EasyQuant | Developed in-house<br>Nat Struct Mol Biol.<br>19:1018-22. PMID:<br>23001004; |  | Used to quantify relative fraction full-length of translated protein from SDS-PAGE |

### Enzymes and chemicals

Enzymes and other reagents were purchased from Thermo Fisher Scientific, New England Biolabs, and Sigma-Aldrich. Oligonucleotides were ordered from Eurofins Genomics. L-[^35^S]-methionine was provided by PerkinElmer. Anti-HA tag antibody (mouse monoclonal) was obtained from BioLegend.

### Cloning and Mutagenesis

The previously described pET-Duet-1 plasmid with N_out_-C_out_ oriented EmrE(C_out_) followed by a variable LepB-derived linker sequence (between 4 and 34 residues), the 9-residue long HA tag, the 17-residue long *E. coli* SecM AP, and a 23-residue long C-terminal tail in MCS2 was used to make all constructs in this study (7, 9). To generate the fused dimer construct the previously described 9TMH-EmrE (C_in_-TMH-C_out_) construct was cloned in place of EmrE(C_out_) in MCS2 of pET-Duet-1 using Gibson assembly^**®**^ (29, 35). For co-expression of EmrE(C_in_) with EmrE(C_out_), the gene encoding the N_in_-C_in_ oriented EmrE(C_in_) version was engineered into MCS1 of pET-Duet-1 harboring EmrE(C_out_) in MCS2 (9). Ordered gene fragments were used to introduce the double mutation G^90^P+G^97^P into EmrE(C_in_). Point mutations in EmrE(C_out_) and deletion of the T7 promoter-2 (32) were done by performing site-specific DNA mutagenesis. The tZ terminator (32) was inserted 25 bp downstream of the EmrE(C_in_) stop codon by Gibson assembly^®^. All cloning and mutagenesis products were confirmed by DNA sequencing. EmrE sequences and the pET-Duet-1 versions used in this study are summarized in Supplementary File S1. The plasmid map in Supplementary Fig. S2 was generated by using SnapGene^®^.

### In vivo pulse-labeling analysis

Induction of protein expression (1 mM IPTG, 10 minutes) followed by [^35^S]-Met pulse-labeling (2 minutes) of BL21 (DE) cells harboring pET-Duet-1 constructs encoding the different EmrE versions and immunoprecipitation using an antibody directed against the HA tag were carried out as previously described (7). In order to detect tag-less EmrE(C_in_) (Supplementary Fig. S2), expression from pET-Duet-1 plasmids carrying EmrE(C_in_) in MCS1 was done using rifampicin to inhibit endogenous transcription (36). In brief, cultures were incubated with 0.2 mg/mL rifampicin after induction and shaken for 15 min at 37°C before radiolabeling. Samples were precipitated, washed and immediately solubilized in SDS sample buffer before incubation with RNase.

Radiolabeled proteins were detected by exposing dried gels to phosphorimaging plates, which were scanned in a Fuji FLA-3000 scanner. Band intensity profiles were obtained using the FIJI (ImageJ) software and quantified with our in-house software EasyQuant. *A*_*c*_ and/or *FL*_*c*_ controls were included in the SDS-PAGE analysis for constructs where the identities of the *A* and *FL* bands were not immediately obvious on the gel. Data was collected from at least three independent biological replicates, and averages and standard errors of the mean (SEM) were calculated. Statistical significance was calculated using a two-sided t-test.

## Acknowledgments

We thank Dr. Gerald Striedner (University of Natural Resources and Life Science, Vienna) for advice on the tZ terminator and Dr. Rickard Hedman (Stockholm University) for programming and maintenance of the EasyQuant software. This work was supported by grants from the Knut and Alice Wallenberg Foundation (2017.0323), the Novo Nordisk Fund (NNF18OC0032828), and the Swedish Research Council (2020-03238) to GvH, and by a Marie Curie Initial Training Network Grant (Horizon 2020, ProteinFactory 642863) to FN.

## Figure legends

**Supplementary File S1**. Amino acid sequences of all constructs.

**Supplementary Table S1**. Experimental *f*_*FL*_ values for all constructs.

**Supplementary Figure S2**. pET-Duet-1 tZ terminator and ΔT7 promoter-2 constructs.

**Supplementary File S1:**
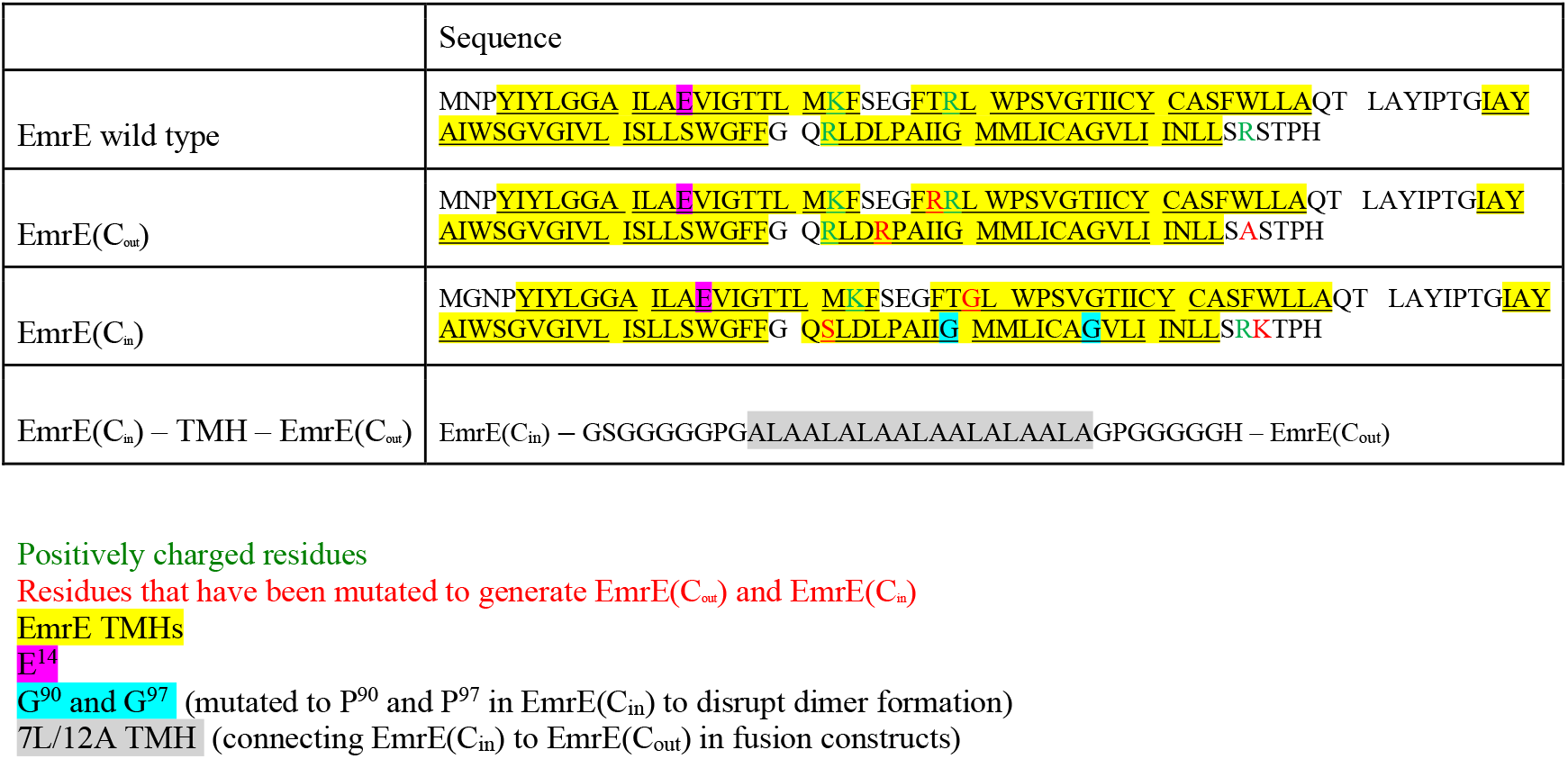
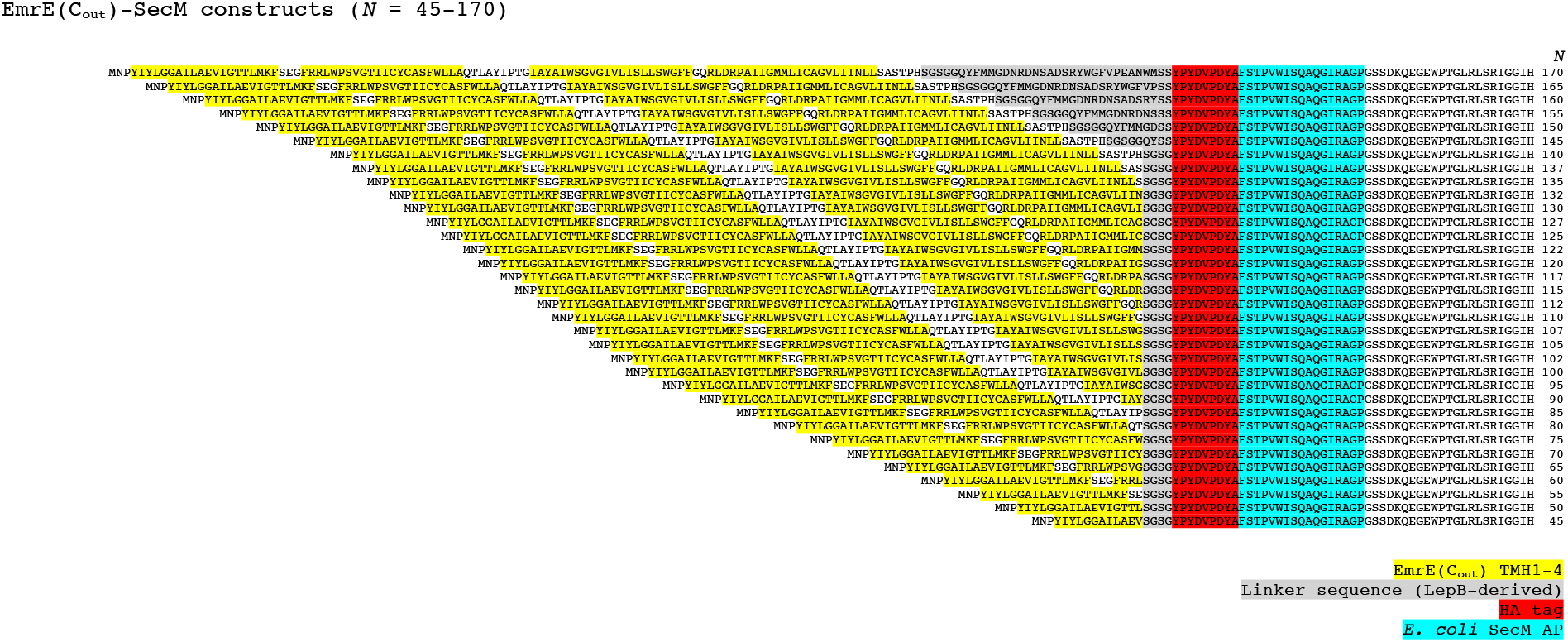
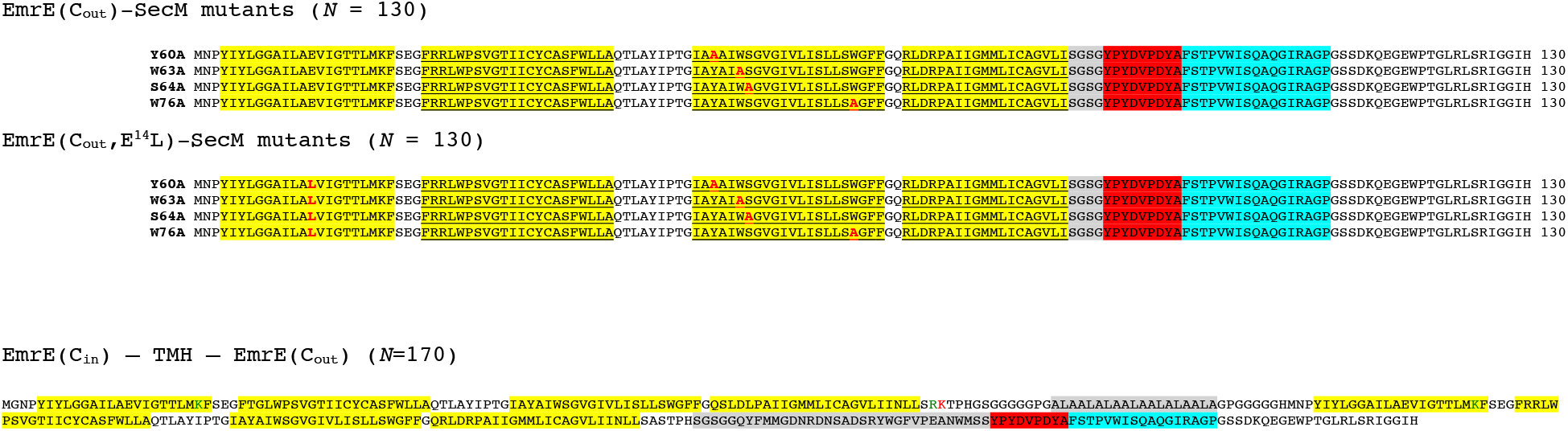
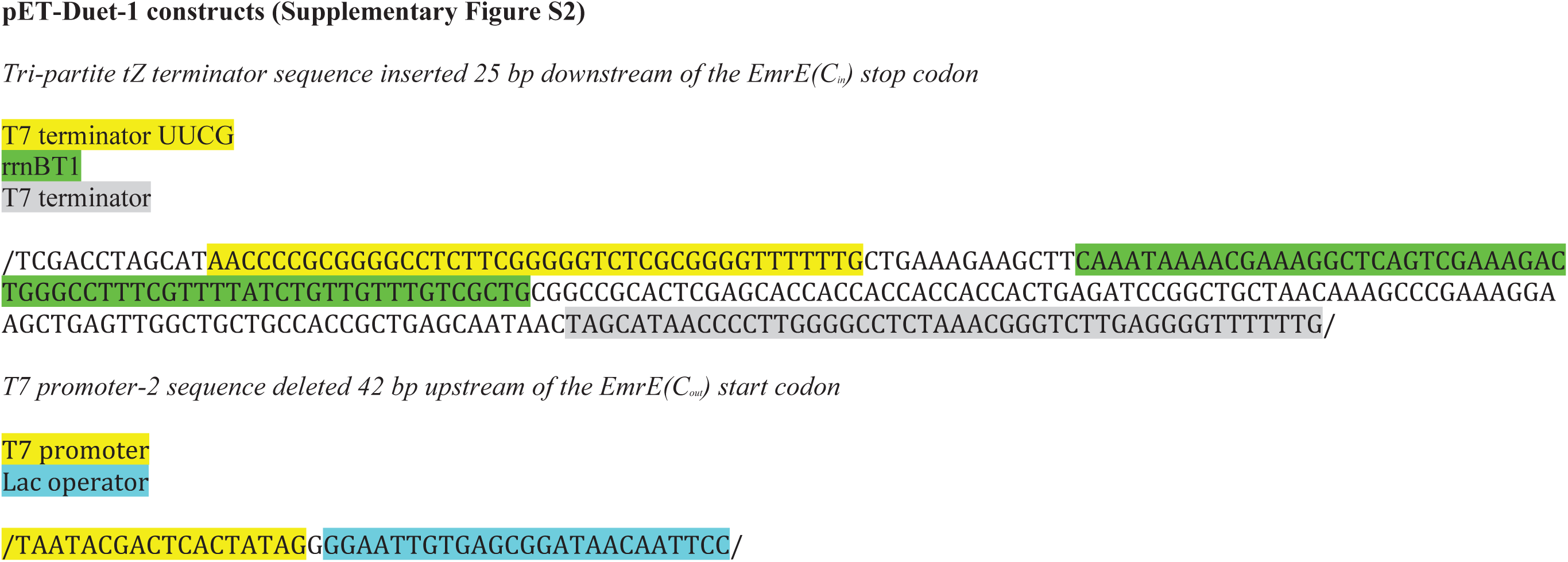
Amino acid sequences of all EmrE constructs plus DNA sequences of pET-Duet-1 constructs.

**Siupplementary Table S1:** Experimental fFL values for all constructs.

**EmrE Cout (Figure 1)**
| N | repeat 1 | repeat 2 | repeat 3 | repeat 4 | repeat 5 | repeat 6 | average | SD | SEM |
| --- | --- | --- | --- | --- | --- | --- | --- | --- | --- |
| 45 | 0.03 | 0.04 |  |  |  |  | 0.03 | 0.01 | 0.01 |
| 50 | 0.06 | 0.12 |  |  |  |  | 0.09 | 0.04 | 0.03 |
| 55 | 0.06 | 0.09 | 0.10 |  |  |  | 0.08 | 0.02 | 0.01 |
| 60 | 0.00 | 0.03 | 0.03 |  |  |  | 0.02 | 0.02 | 0.01 |
| 65 | 0.03 | 0.09 | 0.04 |  |  |  | 0.05 | 0.03 | 0.02 |
| 70 | 0.06 | 0.06 | 0.20 |  |  |  | 0.11 | 0.08 | 0.05 |
| 75 | 0.17 | 0.25 | 0.15 |  |  |  | 0.19 | 0.06 | 0.03 |
| 80 | 0.46 | 0.47 | 0.50 |  |  |  | 0.48 | 0.02 | 0.01 |
| 85 | 0.50 | 0.48 | 0.50 |  |  |  | 0.49 | 0.01 | 0.01 |
| 90 | 0.38 | 0.35 | 0.44 |  |  |  | 0.39 | 0.05 | 0.03 |
| 95 | 0.34 | 0.38 | 0.16 |  |  |  | 0.29 | 0.12 | 0.07 |
| 100 | 0.36 | 0.44 | 0.36 |  |  |  | 0.39 | 0.04 | 0.03 |
| 102 | 0.41 | 0.52 |  |  |  |  | 0.46 | 0.08 | 0.06 |
| 105 | 0.84 | 0.82 | 0.77 |  |  |  | 0.81 | 0.03 | 0.02 |
| 107 | 0.91 | 0.94 | 0.87 |  |  |  | 0.91 | 0.03 | 0.02 |
| 110 | 0.94 | 0.89 | 0.89 |  |  |  | 0.91 | 0.03 | 0.02 |
| 112 | 0.85 | 0.89 | 0.95 |  |  |  | 0.90 | 0.05 | 0.03 |
| 115 | 0.69 | 0.71 | 0.73 |  |  |  | 0.71 | 0.02 | 0.01 |
| 117 | 0.70 | 0.88 | 0.79 |  |  |  | 0.79 | 0.09 | 0.05 |
| 120 | 0.90 | 0.89 | 0.88 |  |  |  | 0.89 | 0.01 | 0.01 |
| 122 | 0.94 | 0.96 | 0.89 |  |  |  | 0.93 | 0.04 | 0.02 |
| 125 | 0.72 | 0.64 | 0.76 |  |  |  | 0.71 | 0.06 | 0.04 |
| 127 | 0.73 | 0.92 | 0.81 |  |  |  | 0.82 | 0.10 | 0.06 |
| 130 | 0.77 | 0.66 | 0.72 | 0.61 | 0.65 | 0.68 | 0.68 | 0.06 | 0.02 |
| 132 | 0.74 | 0.81 | 0.88 |  |  |  | 0.81 | 0.07 | 0.04 |
| 135 | 0.74 | 0.76 | 0.75 |  |  |  | 0.75 | 0.01 | 0.01 |
| 137 | 0.81 | 0.82 | 0.88 |  |  |  | 0.83 | 0.04 | 0.02 |
| 140 | 0.59 | 0.51 | 0.50 |  |  |  | 0.53 | 0.05 | 0.03 |
| 145 | 0.32 | 0.24 | 0.29 |  |  |  | 0.28 | 0.04 | 0.02 |
| 150 | 0.46 | 0.40 | 0.35 |  |  |  | 0.40 | 0.06 | 0.03 |
| 155 | 0.21 | 0.23 | 0.32 |  |  |  | 0.25 | 0.06 | 0.03 |
| 160 | 0.13 | 0.17 | 0.14 |  |  |  | 0.15 | 0.02 | 0.01 |
| 165 | 0.16 | 0.15 | 0.15 |  |  |  | 0.15 | 0.00 | 0.00 |
| 170 | 0.13 | 0.15 | 0.12 |  |  |  | 0.13 | 0.02 | 0.01 |

**EmrE(Cout) [N=130] mutations (Figure 2)**
| background | mutation | repeat 1 | repeat 2 | repeat 3 | repeat 4 | repeat 5 | repeat 6 | average | SD | SEM |
| --- | --- | --- | --- | --- | --- | --- | --- | --- | --- | --- |
| EmrE(Cout) | - | 0.77 | 0.66 | 0.72 | 0.61 | 0.65 | 0.68 | 0.68 | 0.06 | 0.02 |
| EmrE(Cout) | Y60A | 0.66 | 0.74 | 0.65 |  |  |  | 0.68 | 0.05 | 0.03 |
| EmrE(Cout) | W63A | 0.66 | 0.64 | 0.63 |  |  |  | 0.64 | 0.01 | 0.01 |
| EmrE(Cout) | S64A | 0.56 | 0.61 | 0.56 |  |  |  | 0.58 | 0.03 | 0.02 |
| EmrE(Cout) | W76A | 0.63 | 0.63 | 0.62 |  |  |  | 0.63 | 0.01 | 0.00 |
| EmrE(Cout;E14L) | - | 0.54 | 0.57 | 0.64 | 0.52 | 0.54 | 0.51 | 0.55 | 0.05 | 0.02 |
| EmrE(Cout;E14L) | Y60A | 0.67 | 0.63 | 0.60 |  |  |  | 0.63 | 0.04 | 0.02 |
| EmrE(Cout;E14L) | W63A | 0.58 | 0.60 | 0.60 |  |  |  | 0.59 | 0.01 | 0.01 |
| EmrE(Cout;E14L) | S64A | 0.61 | 0.58 | 0.57 |  |  |  | 0.59 | 0.02 | 0.01 |
| EmrE(Cout;E14L) | W76A | 0.63 | 0.56 | 0.61 |  |  |  | 0.60 | 0.04 | 0.02 |

**EmrE(Cin) + EmrE(Cout) (Figure 3)**
| N | repeat 1 | repeat 2 | repeat 3 | average | SD | SEM |
| --- | --- | --- | --- | --- | --- | --- |
| 100 | 0.36 | 0.38 | 0.58 | 0.44 | 0.02 | 0.01 |
| 110 | 0.88 | 0.89 | 0.87 | 0.88 | 0.01 | 0.01 |
| 120 | 0.92 | 0.89 | 0.95 | 0.90 | 0.02 | 0.01 |
| 135 | 0.75 | 0.75 | 0.74 | 0.74 | 0.00 | 0.00 |
| 137 | 0.71 | 0.76 | 0.75 | 0.74 | 0.04 | 0.03 |
| 140 | 0.64 | 0.66 | 0.64 | 0.65 | 0.01 | 0.01 |
| 145 | 0.58 | 0.62 | 0.55 | 0.58 | 0.03 | 0.02 |
| 150 | 0.52 | 0.53 | 0.57 | 0.54 | 0.03 | 0.02 |
| 155 | 0.34 | 0.34 | 0.31 | 0.33 | 0.01 | 0.01 |
| 160 | 0.18 | 0.15 | 0.14 | 0.16 | 0.02 | 0.01 |
| 165 | 0.17 | 0.17 | 0.19 | 0.18 | 0.01 | 0.01 |

**EmrE(Cin-TMH-Cout) (Figure 3)**
| N | repeat 1 | repeat 2 | repeat 3 | average | SD | SEM |
| --- | --- | --- | --- | --- | --- | --- |
| 100 | 0.75 | 0.69 | 0.59 | 0.68 | 0.08 | 0.05 |
| 105 | 0.78 | 0.78 | 0.79 | 0.78 | 0.00 | 0.00 |
| 110 | 0.93 | 0.97 | 0.83 | 0.91 | 0.07 | 0.04 |
| 115 | 0.75 | 0.68 | 0.73 | 0.72 | 0.04 | 0.02 |
| 120 | 0.99 | 0.99 | 0.86 | 0.95 | 0.08 | 0.04 |
| 125 | 0.71 | 0.75 | 0.78 | 0.75 | 0.03 | 0.02 |
| 130 | 0.64 | 0.64 | 0.70 | 0.66 | 0.03 | 0.02 |
| 135 | 0.69 | 0.73 | 0.69 | 0.70 | 0.03 | 0.02 |
| 140 | 0.53 | 0.56 | 0.62 | 0.57 | 0.04 | 0.03 |
| 145 | 0.62 | 0.62 | 0.65 | 0.63 | 0.02 | 0.01 |
| 150 | 0.51 | 0.51 | 0.59 | 0.54 | 0.05 | 0.03 |
| 155 | 0.48 | 0.50 | 0.46 | 0.48 | 0.02 | 0.01 |
| 160 | 0.37 | 0.37 | 0.34 | 0.36 | 0.01 | 0.01 |
| 165 | 0.27 | 0.25 | 0.29 | 0.27 | 0.02 | 0.01 |
| 170 | 0.19 | 0.24 | 0.24 | 0.23 | 0.03 | 0.02 |

**EmrE(Cin; G90P + G97P) + EmrE(Cout) (Figure 3)**
| N | repeat 1 | repeat 2 | repeat 3 | repeat 4 | repeat 5 | average | SD | SEM |
| --- | --- | --- | --- | --- | --- | --- | --- | --- |
| 135 | 0.80 | 0.82 | 0.93 |  |  | 0.85 | 0.07 | 0.04 |
| 137 | 0.77 | 0.70 | 0.74 |  |  | 0.74 | 0.04 | 0.02 |
| 140 | 0.52 | 0.63 | 0.71 |  |  | 0.62 | 0.09 | 0.05 |
| 145 | 0.45 | 0.48 | 0.52 | 0.44 | 0.46 | 0.47 | 0.03 | 0.01 |
| 150 | 0.43 | 0.44 | 0.46 |  |  | 0.44 | 0.02 | 0.01 |
| 155 | 0.24 | 0.34 | 0.32 |  |  | 0.30 | 0.05 | 0.03 |
| 160 | 0.12 | 0.24 | 0.16 |  |  | 0.17 | 0.06 | 0.03 |
| 165 | 0.17 | 0.24 | 0.27 |  |  | 0.23 | 0.05 | 0.03 |

**EmrE(Cin-TMH-Cout; G90P + G97P) (Figure 3)**
| N | repeat 1 | repeat 2 | repeat 3 | average | SD | SEM |
| --- | --- | --- | --- | --- | --- | --- |
| 135 | 0.71 | 0.71 | 0.73 | 0.72 | 0.01 | 0.01 |
| 140 | 0.58 | 0.58 | 0.57 | 0.58 | 0.01 | 0.00 |
| 145 | 0.46 | 0.45 | 0.43 | 0.45 | 0.02 | 0.01 |
| 150 | 0.57 | 0.55 | 0.52 | 0.55 | 0.02 | 0.01 |
| 155 | 0.48 | 0.37 | 0.45 | 0.44 | 0.06 | 0.03 |
| 160 | 0.22 | 0.21 | 0.20 | 0.21 | 0.01 | 0.01 |
| 165 | 0.26 | 0.20 | 0.23 | 0.23 | 0.03 | 0.02 |
| 170 | 0.31 | 0.20 | 0.28 | 0.27 | 0.05 | 0.03 |

**EmrE(Cin) + EmrE(Cout) expressed from pET-Duet 1 with the T7 promoter-2 deleted (Supplementary Figure S2)**
| N | repeat 1 | repeat 2 | repeat 3 | average | SD | SEM |
| --- | --- | --- | --- | --- | --- | --- |
| 135 | 0.75 | 0.63 | 0.79 | 0.72 | 0.08 | 0.05 |
| 140 | 0.56 | 0.58 | 0.63 | 0.59 | 0.03 | 0.02 |
| 145 | 0.55 | 0.61 | 0.55 | 0.57 | 0.04 | 0.02 |
| 150 | 0.47 | 0.53 | 0.55 | 0.52 | 0.04 | 0.02 |
| 155 | 0.32 | 0.30 | 0.36 | 0.33 | 0.03 | 0.02 |
| 160 | 0.16 | 0.19 | 0.12 | 0.15 | 0.03 | 0.02 |
| 165 | 0.22 | 0.18 | 0.21 | 0.21 | 0.02 | 0.01 |

**EmrE(Cin) + EmrE(Cout) expressed from pET-Duet 1 with the Tz terminator inserted (Supplementary Figure S2)**
| N | repeat 1 | repeat 2 | repeat 3 | average | SD | SEM |
| --- | --- | --- | --- | --- | --- | --- |
| 135 | 0.67 | 0.77 | 0.68 | 0.71 | 0.06 | 0.03 |
| 137 | 0.69 | 0.62 | 0.65 | 0.66 | 0.03 | 0.02 |
| 140 | 0.54 | 0.55 | 0.52 | 0.54 | 0.01 | 0.01 |
| 145 | 0.59 | 0.56 | 0.63 | 0.60 | 0.04 | 0.02 |
| 150 | 0.56 | 0.54 | 0.55 | 0.55 | 0.01 | 0.00 |
| 155 | 0.40 | 0.38 | 0.34 | 0.37 | 0.03 | 0.02 |
| 160 | 0.15 | 0.18 | 0.23 | 0.19 | 0.04 | 0.02 |
| 165 | 0.19 | 0.19 | 0.22 | 0.20 | 0.01 | 0.01 |

**Supplementary Figure S1.**
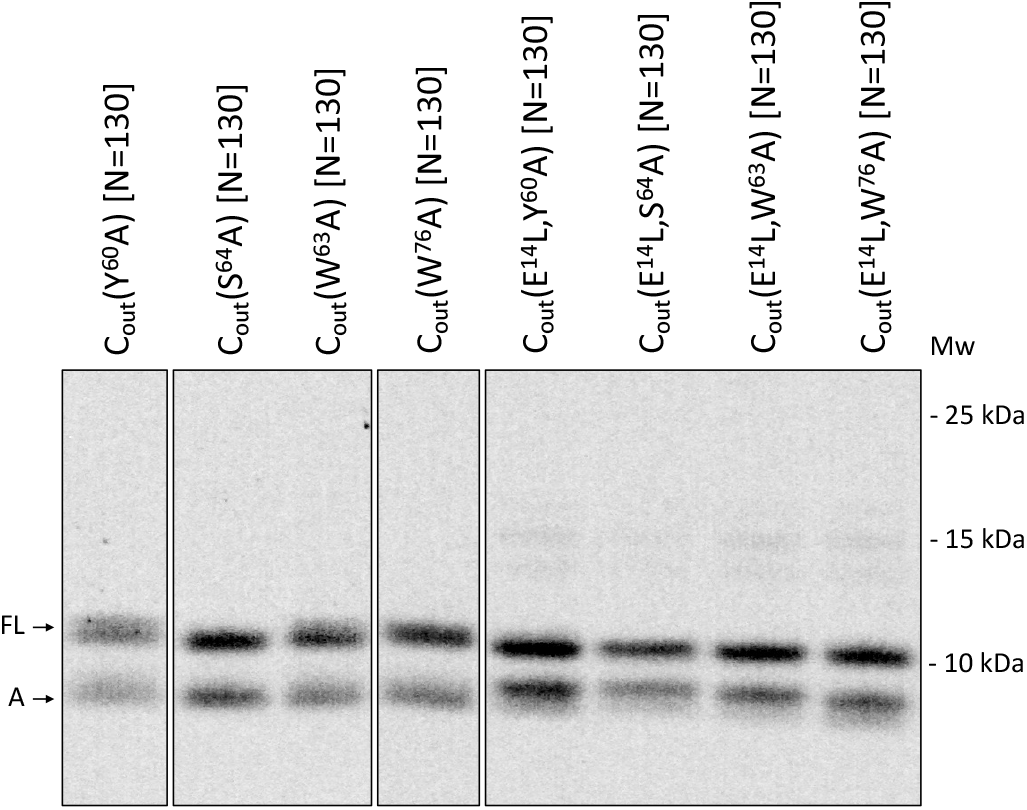
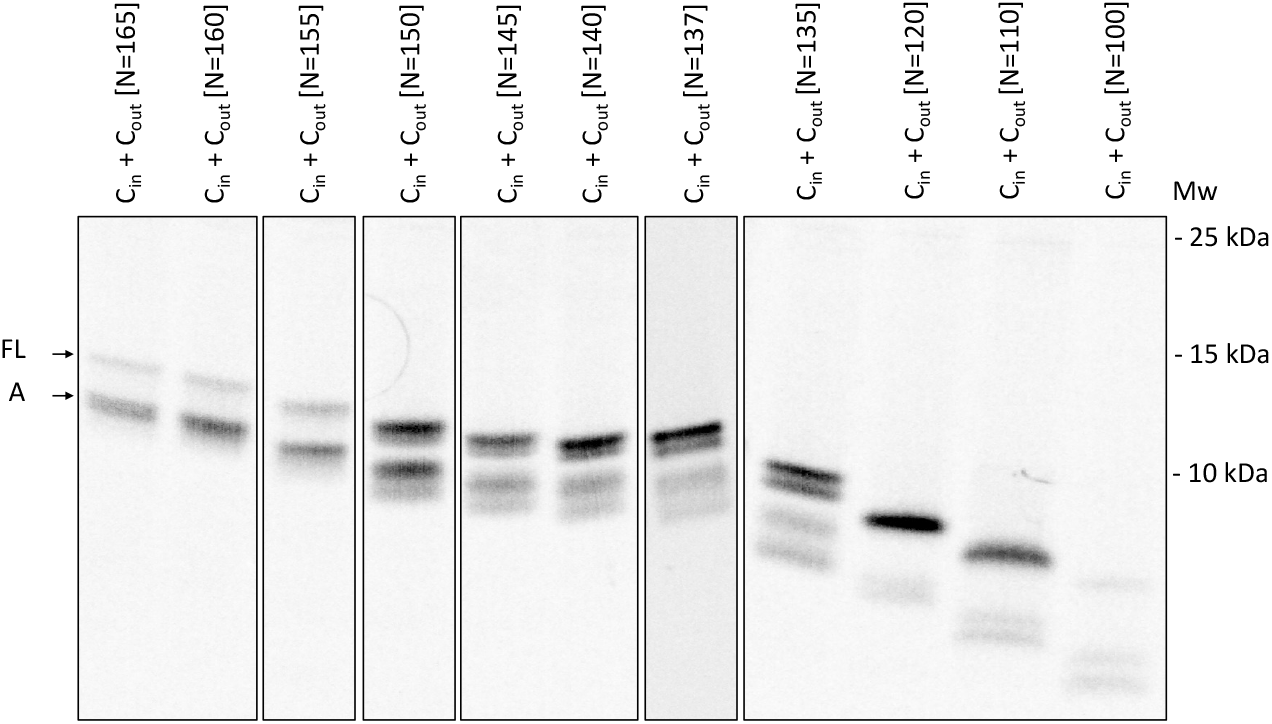
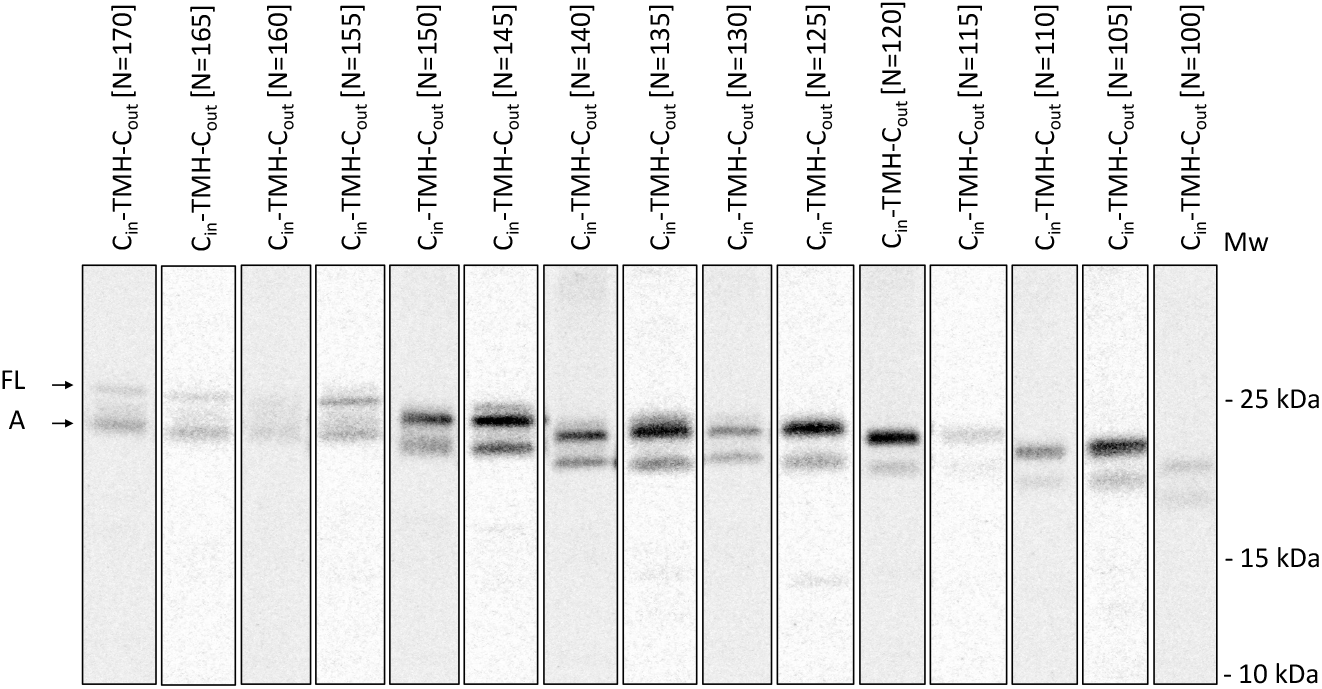
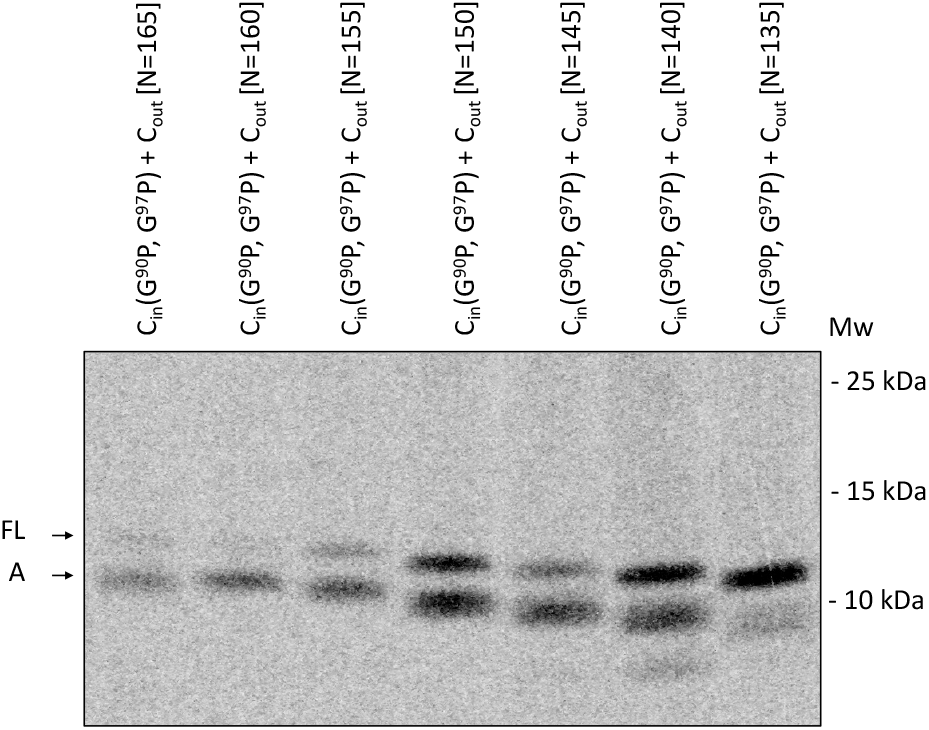
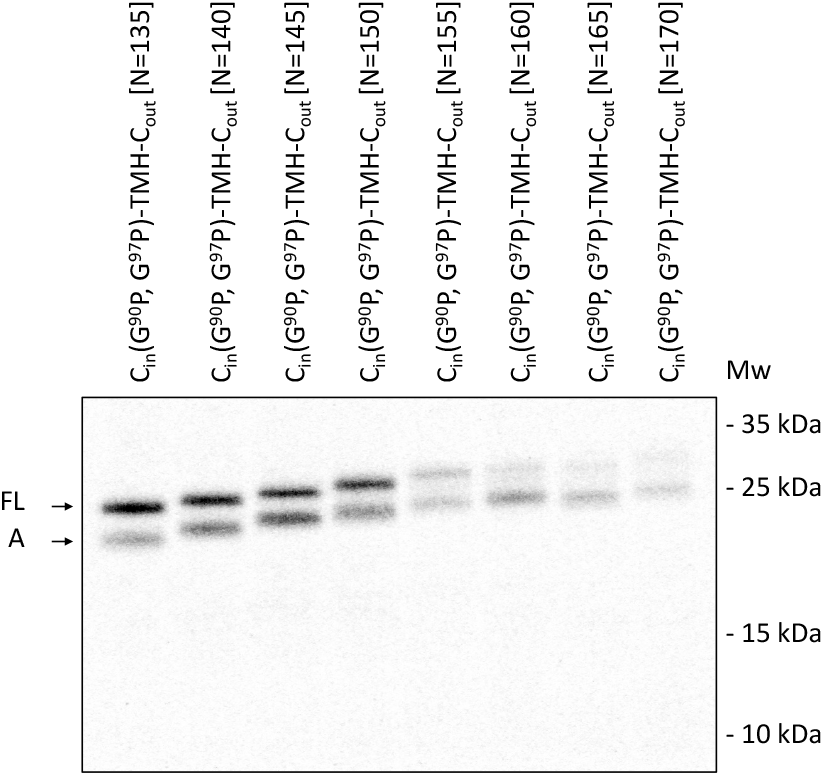
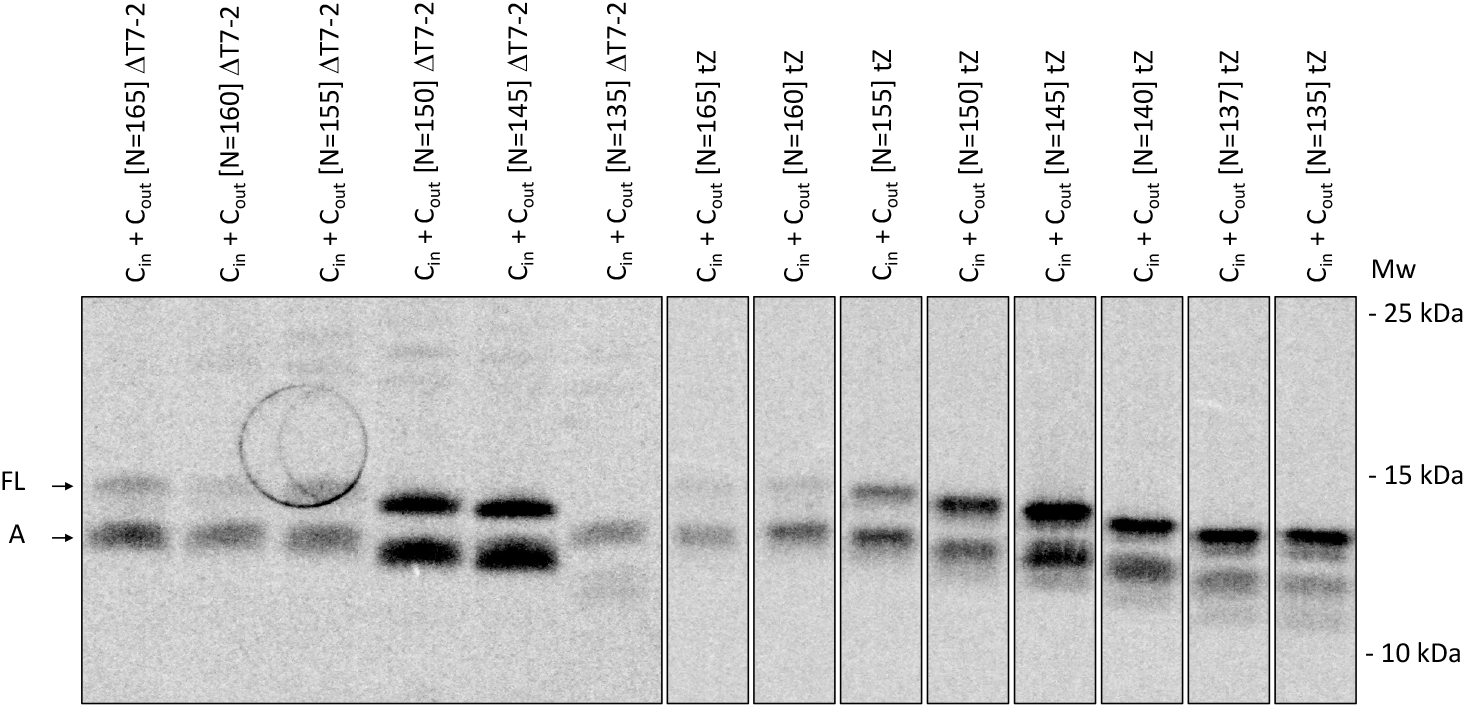
SDS-PAGE gels for all constructs.

**Supplementary Figure S2.**
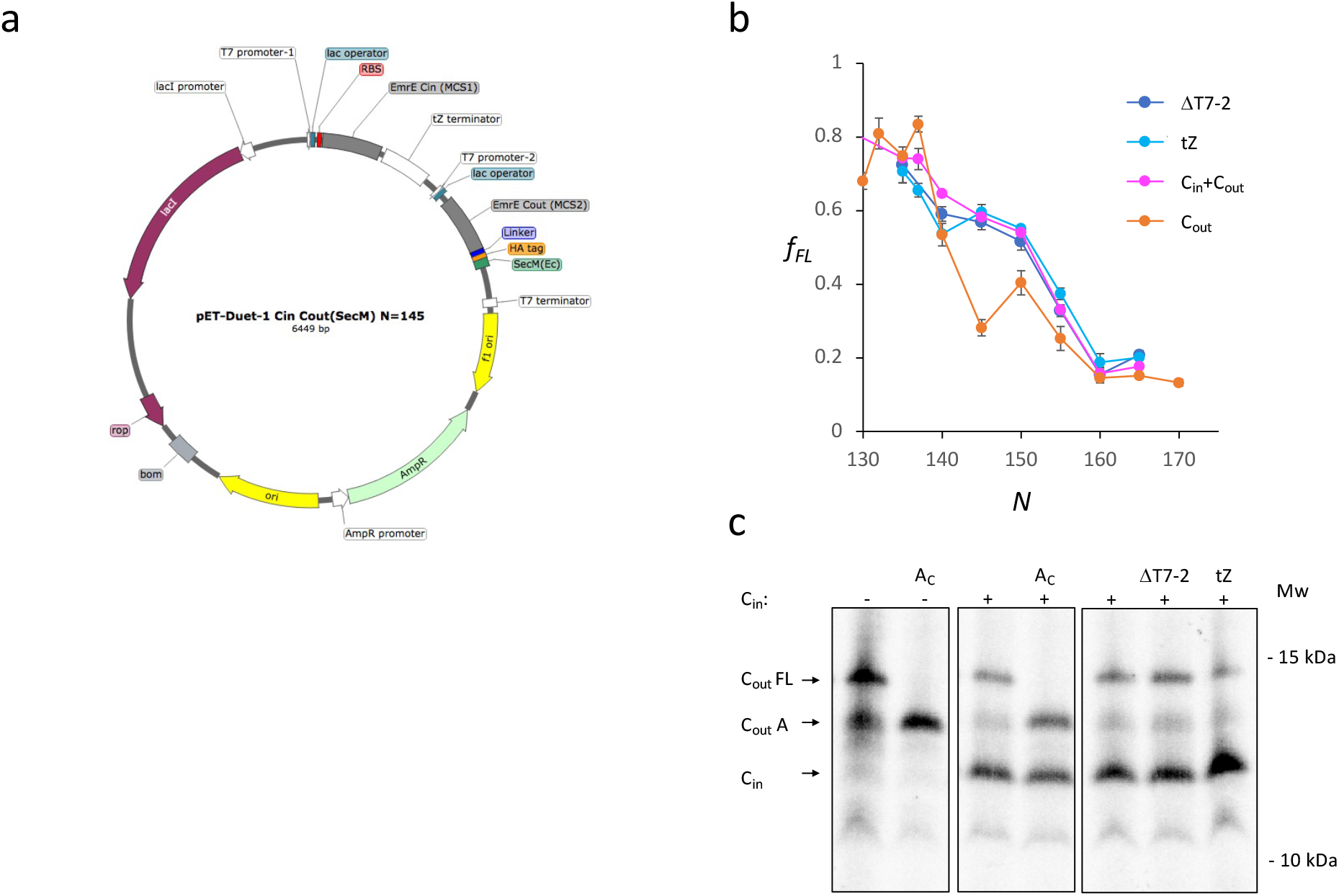
**(a)** pET-Duet-1 vector used for co-expression of EmrE(C_in_) with EmrE(C_out_) constructs. Note the tZ terminator insertion and T7 promoter-2 deletion used for the expressions shown in panels *b* and *c*. **(b)** FPs for EmrE(C_out_) (orange data points), EmrE(C_out_) with co-expressed EmrE(C_in_) (magenta data points), EmrE(C_out_) with co-expressed EmrE(C_in_) from pET-DUET-1 with the T7 promoter-2 deleted (dark blue data points), and EmrE(C_out_) with co-expressed EmrE(C_in_) from pET-Duet-1 with the tZ terminator inserted between the EmrE(C_in_) and EmrE(C_out_) ORFs (light blue data points). **(c)** SDS-PAGE gel showing [^35^S]-Met labelled EmrE(C_in_) and EmrE(C_out_)-SecM [*N*=145] products. To inhibit endogenous transcription, constructs were expressed in the presence of rifampicin. Lane 1 shows EmrE(C_out_)-SecM [*N*=145] expressed in the absence of EmrE(C_in_) and lane 2 shows the corresponding arrest control. Lane 3 shows the coexpression of EmrE(C_in_) with EmrE(C_out_)-SecM [*N*=145] and lane 4 shows the corresponding arrest control. Lane 5 is a repeat of lane 3. Lane 6 shows EmrE(C_out_)-SecM [*N*=145] co-expressed with EmrE(C_in_) from pET-Duet-1 with the T7 promoter-2 deleted. Lane 7 shows EmrE(C_out_)-SecM [*N*=145] co-expressed with EmrE(C_in_) from pET-Duet-1 with the tZ terminator inserted between the EmrE(C_in_) and EmrE(C_out_)-SecM ORFs.

